# Social dynamics of vervet monkeys are dependent upon group identity

**DOI:** 10.1101/2023.06.02.543415

**Authors:** Elena Kerjean, Erica van de Waal, Charlotte Canteloup

**Author notes:** These authors have contributed equally to this work and share last authorship (Erica van de Waal, Charlotte Canteloup).

## Abstract

Traditions are widespread across the animal realm. Here, we investigated inter-group variability of social dynamics in wild vervet monkeys (*Chlorocebus pygerythrus*). We analysed 84 704 social behavioural interactions involving 247 individuals collected over nine years in three neighbouring groups of wild vervet monkeys. We found that, in one group - Ankhase - individuals had a higher propensity to be affiliative (i.e., sociality) and grooming interactions were more reciprocal. Despite yearly fluctuations in sociality, differences between groups remained stable over time. Moreover, our statistical model predictions confirmed that these findings were maintained for similar sex ratios, age distributions and group sizes. Strikingly, our results suggested that dispersing males adapted their sociality to the sociality of the group they integrated with. As a whole, our study sheds light on the existence of stable social dynamics dependent upon group identity in wild vervet monkeys and suggests that at least part of this variability is socially mediated.

**Graphical Abstract:** 

**Highlights:**

- The sociality of vervet monkeys groups differs consistently across a nine years study despite similar genetic and ecological environments.
- Dispersing males adapt their sociality according to the group they integrate with.
- In the most social group, grooming interactions were more reciprocal.

## 1. INTRODUCTION

Across time and space, culture shapes us: from what we know, to how we behave and interact - it fuels the diversity of human behaviours. Culture encompasses “those group-typical behavior patterns shared by members of a community that rely on socially learned and transmitted information”^1^ and includes several traditions - which are defined as “a behavioral practice that is relatively enduring (i.e., is performed repeatedly over a period of time), that is shared among two or more group members, and that depends in part on socially aided learning for its generation in new practitioners”^2^. The accumulation of evidence in wild and captive non-human animals (hereafter: animals) revealed the existence of traditions in a variety of mammals and birds, from tool use in chimpanzees to song dialects in birds^3^. Although traditions appear to be widespread in non-human animals, the use of the term “culture” for non-humans is still debated. While, some researchers consider that culture relies on complex cognitive processes involving symbols and that cumulative culture is a hallmark of human beings^4,5^, depending on know-how copying processes^6^, others adopt a broader definition of culture defined as socially transmitted knowledge which is not based on genetic inheritance^7,8^. It has been proposed that the difference between human and animal culture might rely on the fact that, contrary to other animals, humans are able to generate knowledge that accumulate over generations ^9,5^.

Over the last decades, animal tradition studies not only helped researchers to better understand the diversity of intra-species behavioural repertoires, but also to unravel the evolutionary roots and the selective pressures leading to the emergence of culture. A pioneer study reported the innovation and spread of sweet potato washing in a group of Japanese macaques (*Macaca fuscata*) on Koshima Islet, Japan^10^. Such behaviour was observed only in a few groups of Japanese macaques^4^, authors then described this behaviour as pre-cultural^10^. However, besides observing intra-species behavioural variability between groups (i.e., a social unit of individuals) or between populations (i.e., several groups within a geographically defined area), a fundamental aspect of culture is that cultural behaviours should be transmitted through social learning, where social learning is defined as a learning process “facilitated by observation of, or interaction with, another individual (or its products)”^11^. In this example, Koshima group was the only one where the sweet potato washing behaviour progressively spread : by 1956, eleven monkeys of the same group acquired this behaviour (80 % of adults). What remains controversial in this example is the mechanism implied in the spread of the behaviour; imitation being unlikely to explain the adoption of the habit that could be due to simpler mechanisms (e.g. enhancement; social facilitation)^4^. A classical method used to investigate the social origin of a behaviour is to exclude both obvious ecological and genetic variations, leaving social transmission as the only probable cause of the observed behavioural diversity (i.e., method of exclusion)^12^. However, across the literature, most studies have investigated the behavioural variability of geographically distant populations^3^. Quantifying the influence of ecological and genetic variation in such a setting can easily be misleading in the case of subtle environmental variations^13^. Recently, several researchers argued that studying neighbouring groups of animals of the same species limits genetic and ecological biases because they live in relatively similar ecological and genetical environments^14,15^. Thus, what was before a neglected scale of behavioural diversity - the inter-group level - is now resurgent in recent literature and offers significant advantages in the study of traditions in wild animals ^14,16,17,18,19,20^.

Ecological and genetic environments are however not the only source of variability which should be controlled for: group size, sex ratio or age distribution - broadly defined as group demographic variables - are another stream of variability which could explain inter-group behavioural diversity^21^. In species with sex or age specific behaviours, slight differences in proportions of the sex ratio and/or the age distribution could lead to specific group behavioural patterns. For example, in vervet monkeys, females spend more time grooming conspecifics than males^22,23^ and thus a group with a higher proportion of females may overall dedicate more time to grooming rather than a group with a balanced sex ratio. DeTroy et al.^24^, recently highlighted the influence of group demographics on co-feeding tolerance in four groups of sanctuary-housed chimpanzees across eight years. They found that co-feeding tolerance increased with the number of juvenile females and decreased as the number of infants increased. This study stressed that long-term studies have the potential to quantify and take into account group demographic variation across time and can extend our understanding of animal traditions, especially in animals with slow life histories such as primates ^25,24^**^?^**.

Animal traditions can be classified in three different categories: material, food-related and social traditions (see^26^ for a review of monkey traditions). Even though, several social behaviours are known to be transmitted socially^27,28,29^, strong evidence for social traditions is still lacking in comparison to material or food-related animal traditions^30^. Some studies even suggested a relative higher importance (41%) of social traditions over other types of traditions in specific species such as spider monkeys (*Ateles geoffroyi*)^31^. Qualitative (presence/absence) and quantitative (frequency) differences in non-human primates’ (hereafter: primates) social behaviour have been reported. In comparison to qualitative differences in behaviour, quantitative behavioural diversity in primate social behaviour has been less investigated so far. However, when investigating the variation in the frequency of wild spider monkey behaviours (e.g. foraging, greetings and resting), Santorelli et al.^32^ found a significant amount of variation and identified them as “candidate traditions”. Quantitative variations in inter-species interactions have also been reported in different species of monkeys (*Cebus capucinus*, *Macaca fascicularis*)^33,34,35^. For example, over a four-year period, Rose et al.^33^ found that one group of white-faced capuchin monkeys (*Cebus capucinus*) was highly involved in grooming with spider monkeys compared to five other neighbouring groups. De Waal & Johanowicz^36^ reported that juvenile rhesus macaques (*Macaca mulatta*) dramatically increased their reconciliation rates after being translocated to a group of a more conciliatory species, the stumptail macaques (*Macaca arctoides*). In 2004, Sapolsky et al.^16,37^ described the emergence and persistence of what they called a “pacific culture” in a group of wild olive baboons (*Papio anubis*). In the mid-1980s, half of the males from their study group died from tuberculosis. Two decades later, all males from the original study died and were replaced by immigrants, and no adult males were native to the group anymore. Researchers observed a significantly higher inter-sexual grooming rate and reduced spatial proximity, as well as a “relaxed” hierarchy between males where low-rankers had a higher probability of accessing contested resources. Incoming males first showed more usual aggressive patterns before conforming to the group social specificity. In the above-cited studies, such global changes in social behaviours were described as “social style”, “social norms”, “social climates”, or “social dynamics”^38^. Here, we use the term “social dynamics” to align with research on great ape social traditions^39^. Overall, even if quantitative variability in social behaviours have been identified, more evidence that these differences in social dynamics are mainly dependent upon social learning in monkeys is still lacking.

Primate social behaviours are classified either as affiliative (friendly) or agonistic (aggressive and submissive). No broad agreement exists on how to measure group sociality. It can be assessed via social tolerance metrics (co-feeding and spatial proximity networks) or pro-social indices (food sharing indices)^25,24,40^, but a variety of other metrics have been used in the literature^41,42^. Based on the definition of sociality in chimpanzees as “interaction tendencies”^39^, we will use the term “sociality” to refer to the propensity to be affiliative (i.e., the relative amount of affiliative interactions compared to agonistic ones) either at the individual level or at the group level. Among affiliative behaviours, grooming is the most widespread and common in primates^43^. It has a crucial role in primate sociality: it maintains social bonds between individuals and decreases stress^43^. The definition of reciprocity in grooming behaviours (i.e., after A groomed B, the probability of B to groom A increases) lacks a consensus concerning the time-lapse in which reciprocity occurs. Here we align with the recent definition of O’Hearn et. al.,^44^ of a global long-term reciprocity (hereafter: reciprocity) which stands for grooming behaviours that can be exchanged over timescales ranging from hours to years.

In the present study, we aim to discover the differences in social dynamics between three neighbouring groups of wild vervet monkeys (*Chlorocebus pygerythrus*) over nine years and to understand if these differences could be interpreted as different social traditions between groups.

Vervet monkeys are a well-suited species to investigate social traditions because they live in multi-male, multi-female groups, providing individuals with many opportunities to socially learn from different individuals^45^. A body of studies has shown that vervet monkeys socially learn how to manipulate artificial fruits^46,47,14,48,49^ and to process novel food^50^, but their potential to socially learn global social dynamics remains poorly known. Here, we focused on two main aspects of primate social life. First, sociality (i.e., propensity to be affiliative) and second, grooming reciprocity (i.e., how reciprocal are grooming exchanges). We asked three main questions: (I) Is sociality influenced by group identity across years? If yes, do dispersing males adapt their sociality when they integrate into a new group? (II) Is sociality similarly distributed between individuals in each group according to their socio-demographic category and kinship? (III) Are differences in grooming reciprocity between groups consistent with differences in sociality? Here, we extend previous studies by combining three grounding aspects of research on animal traditions: the study of several neighbouring groups with overlapping home-ranges and migration flux; the quantification of the influence of group demographics, and the use of a continuous long term database. We used nine years of observational data from three neighbouring groups of wild vervet monkeys from the Inkawu Vervet Project (IVP) : Ankhase (AK), Baie Dankie (BD) and Noha (NH). Borgeaud et al.,^51^ suggested in 2016 that the stability and structure of vervet monkey grooming networks differed over a two-year period using a social network-based analysis. Previous studies also found inter-group differences in co-feeding tolerance between adult females of AK, BD, and NH^52^. However, these studies did not investigate the long-term stability of these differences nor the influence of group demographical conditions. In the later study^52^, one of our study groups, AK, displayed a higher level of co-feeding tolerance. Based on these previous findings and regarding question I, we predict AK to have an overall higher propensity to be affiliative than NH and BD. Additionally, van de Waal et al.^47^ showed that dispersing males seem to conform to local norms in their food choice, abandoning their natal preference. Thus, regarding question I, we predict that dispersing males might also conform to the local sociality. Concerning question II, we predict that individuals from all socio-demographic categories in AK are more social than individuals in the respective socio-demographic categories in BD and NH, and the effect of kinship to be similar across groups. Previous analysis also suggested that females of distant ranks more regularly groom each other in AK^51^. Thus, regarding question III, we predict grooming reciprocity to be also higher in AK. If such differences are stable over years and independent from group demography, it will favor an effect of group identity on social dynamics.

## RESULTS

### Question I: Groups’ sociality differs congruently across years

Model 1.0 revealed that sociality was significantly different between the three groups of vervet monkeys and that this difference was stable across time (Figure 1A, Table 1). Indeed, *Group* had a significant effect on sociality (Likelihood ratio-test *LRT_Group_* = 729.47, *p <* 0.001) whatever the year. In particular, AK had a significantly higher sociality than BD (Average marginal effect *AME_AK−BD_* = 0.06, *p <* 0.001) and NH (*AME_AK−NH_* = 0.1, *p <* 0.001), and BD had a significantly higher sociality than NH (*AME_NH−BD_* = 0.03, *p <* 0.001). Furthermore, according to Model 1.1 the sociality of each group increased with time. AK and BD followed a similar trend (Conditional mode *CM_Y ears|AK_* = 0.17; *CM_Y ears|BD_* = 0.16) while NH trend was slightly slower (*CM_Y ears|BD_* = 0.09). These statistical results are illustrated by the quasi-constant superiority of AK sociality and the relatively parallel and positive evolution of groups’ sociality over years in each group (Figure 1A).

**Figure 1.**
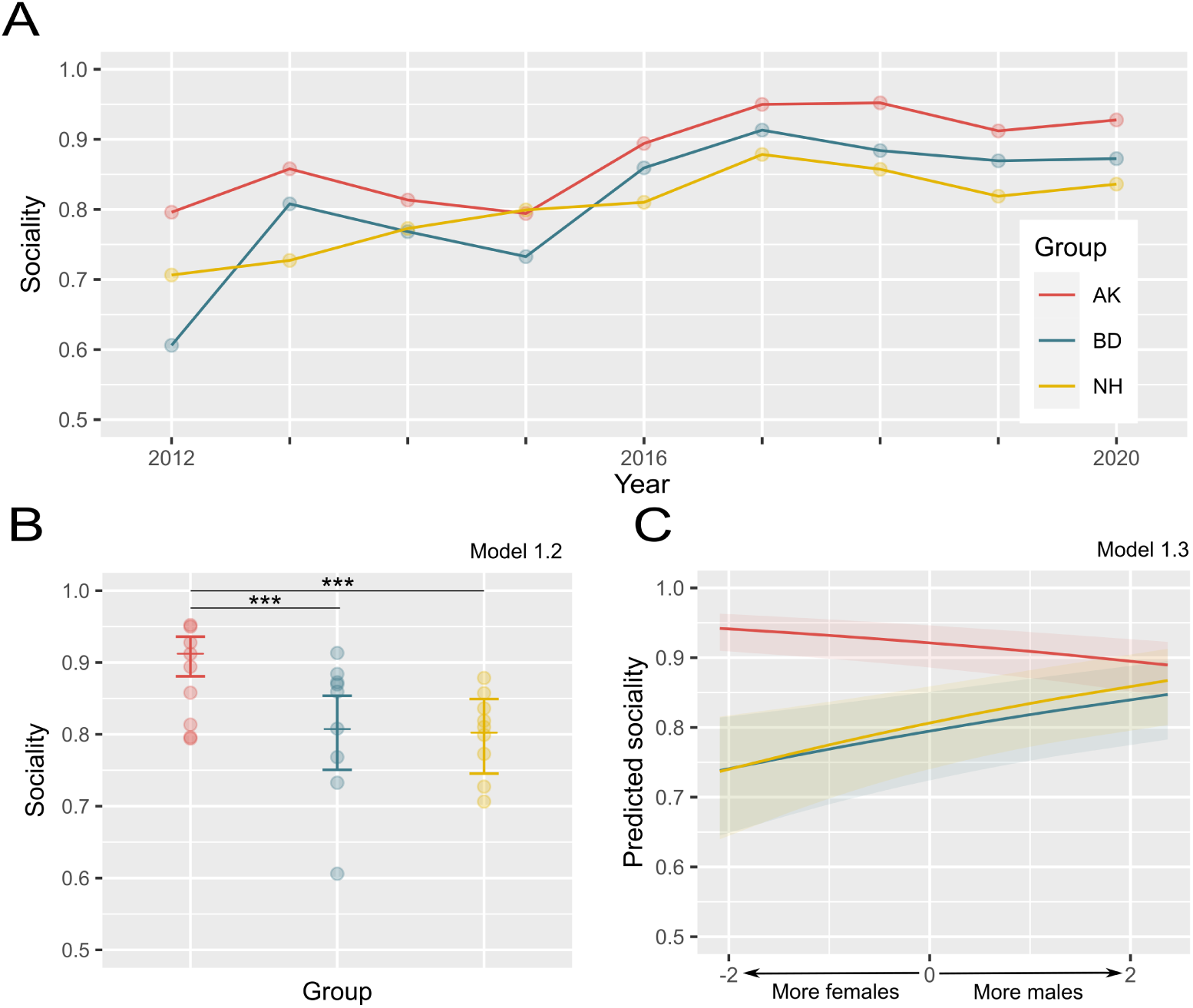
Groups’ sociality depends on group identity. A) Dynamics of the sociality of AK, BD and NH across years. Dots represent the sociality of each group for every year of the study as computed from raw data. B) Predicted sociality of AK, BD and NH in similar demographic environments. The small horizontal bars denote the predicted mean of sociality within each group and the error bars denote the standard error of the mean (SEM) as predicted by Model 1.2. Dots represent the sociality of each group for every year of the study as computed from raw data. (***) denotes the significant differences at the *α <* 0.001 level. C) The effect of group identity as a function of the different values of sex ratio within the groups. The x-axis represents the variation in the scaled value of the sex ratio, negative values represent a female skewed group and positive values represent a male skewed group compared to the mean observed sex ratio. 95% confidence interval (CI). Graphical representations are based on estimates from Model 1.3. AK: Ankhase, BD: Baie Dankie, NH: Noha.

**Table 1.**
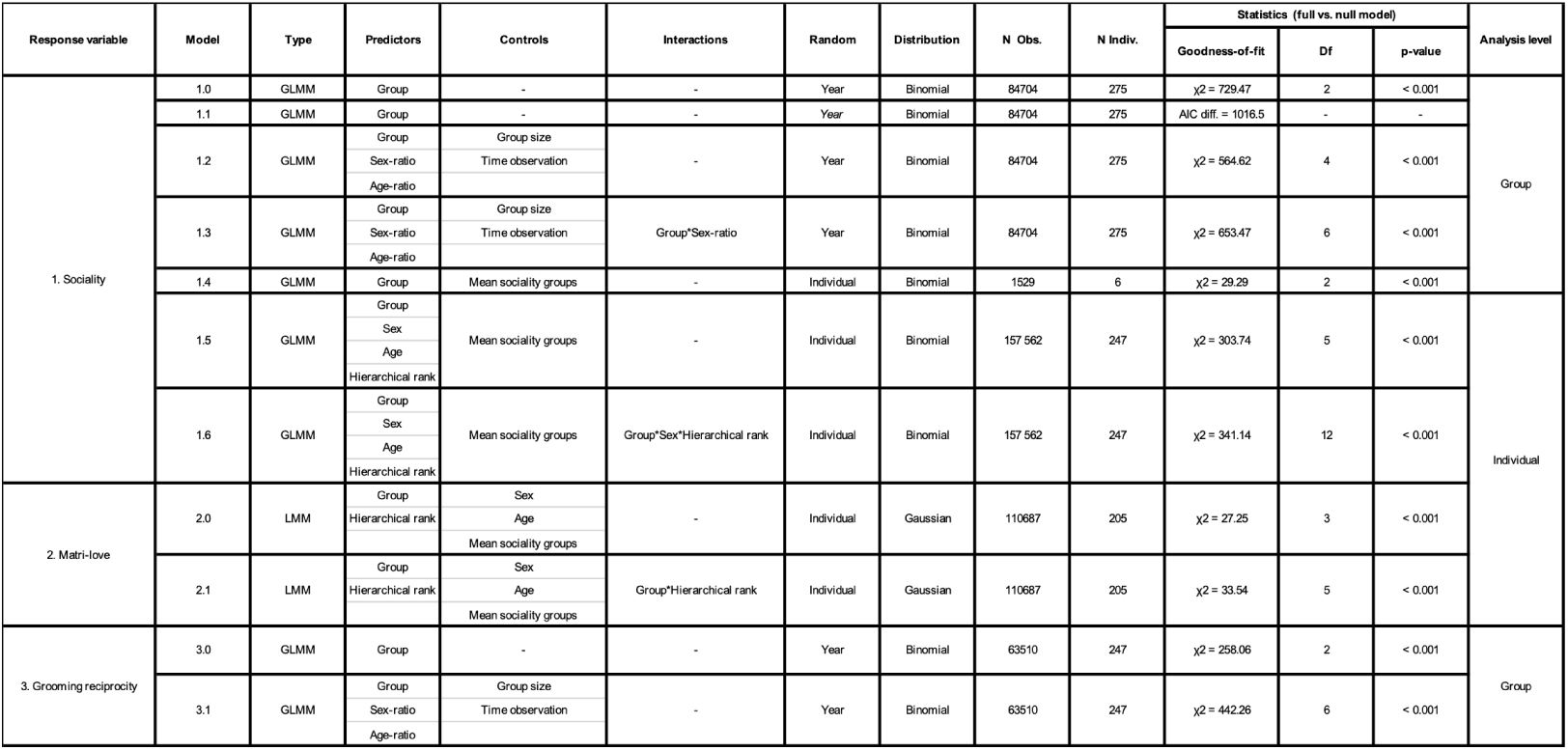
Statistical models summary.: P-values correspond to the full vs. null model comparisons for each model. Variables in *italic* in the random column correspond to random slopes while the one not in italic correspond to random intercepts. Dev.: Deviance, Df: Degree of freedom, diff. : difference, GLM: Generalized Linear Model, GLMM: Generalized Linear Mixed-Model, Indiv : Individual, LMM: Linear Mixed Model, Obs.: Observation.

### Question I: AK has a higher sociality than other groups in similar demographic environments

Model 1.2 predicted that for similar age ratios, sex ratios and group sizes, group identity still had a significant effect on sociality (*LRT_Group_*= 563.01, *p <* 0.001) (Figure 1B, Table 1). In such a setting, AK sociality was predicted to be 11% higher than BD sociality and 12% higher than NH sociality (*AME_AK−BD_* = 0.11, *p <* 0.001; *AME_AK−NH_* = 0.12, *p <* 0.001). However, in contrast to Model 1.0, we found no significant difference between the sociality of BD and of NH (*p* = 1.00). Regarding the global effect of demography on groups’ sociality, we found that sociality decreased as the relative number of males or adults increased in the group (*AME_Age−ratio_* = *−*0.008, *p* = 0.001; *AME_Sex−ratio_* = *−*0.01, *p <* 0.001).

### Question I: When all groups contain more females, differences in sociality are stronger

At a finer scale, we found evidence that the effect of group identity on sociality was stronger when all groups included more females than males (i.e., low sex ratio) (Figure 1C). Indeed, we found that the sex ratio significantly modified the influence of group identity on sociality (*LRT_Group∗Sex−ratio_* = 88.85, *p <* 0.001). The results of Model 1.3 (Table 1) revealed that in AK, a relative increase of females induced a significant increase of sociality (*β_Sex−ratio_* = *−*0.16, *p <* 0.001) while the opposite effect was observed in BD (*β_Sex−ratio_* = 0.15, *p* = 0.001) and NH (*β_Sex−ratio_* = 0.19, *p* = 0.002). As a consequence, when all groups included relatively more females (*Sex ratio*: *−*2.09), Model 1.3 predicted that the sociality of the different groups diverged, AK being significantly more social than the two other groups (*AK > BD ∼ NH*: *β_BD−AK_* = *−*1.75, *p <* 0.001; *β_NH−AK_* = *−*1.76, *p <* 0.001; *BD − NH*: *p* = 1.00). In contrast, when relatively more males were present in the group (*Sex ratio*: 2.3658) differences faded away. In such a setting, our model predicted that only AK and BD sociality were significantly different from each other (*AK > BD*: *β_BD−AK_* = *−*0.37, *p* = 0.004) but neither AK nor BD was different from NH (*AK − NH*: *p* = 1.00, *NH − BD*: *p* = 1.00). Furthermore, within every group a mild interaction between *Sex ratio*\**Age-ratio* was detected (*LRT_Sex−ratio∗Age−ratio_* = 42.29, *p <* 0.001; see Figure S1).

### Question I: Adult males become less social when they disperse to BD and NH compared to AK

Comparing the sociality of dispersing males before and after they dispersed (during their first year of dispersal) between our study groups revealed that their sociality was on average higher by +0.13 *±* 0.12 (*N* = 5, Figure 2 and Figure 3) when they integrated into AK rather than when they dispersed to BD or NH. Six adult males dispersed within our three study groups during our study (Figure 2). Four males dispersed between AK and NH (WOL, TWE, YAN, KEK), one male dispersed from AK to BD (HAM), and one male dispersed from NH to BD (CHE). Only two of those males stayed a second year in both of their dispersing groups (CHE, HAM).

**Figure 2.**
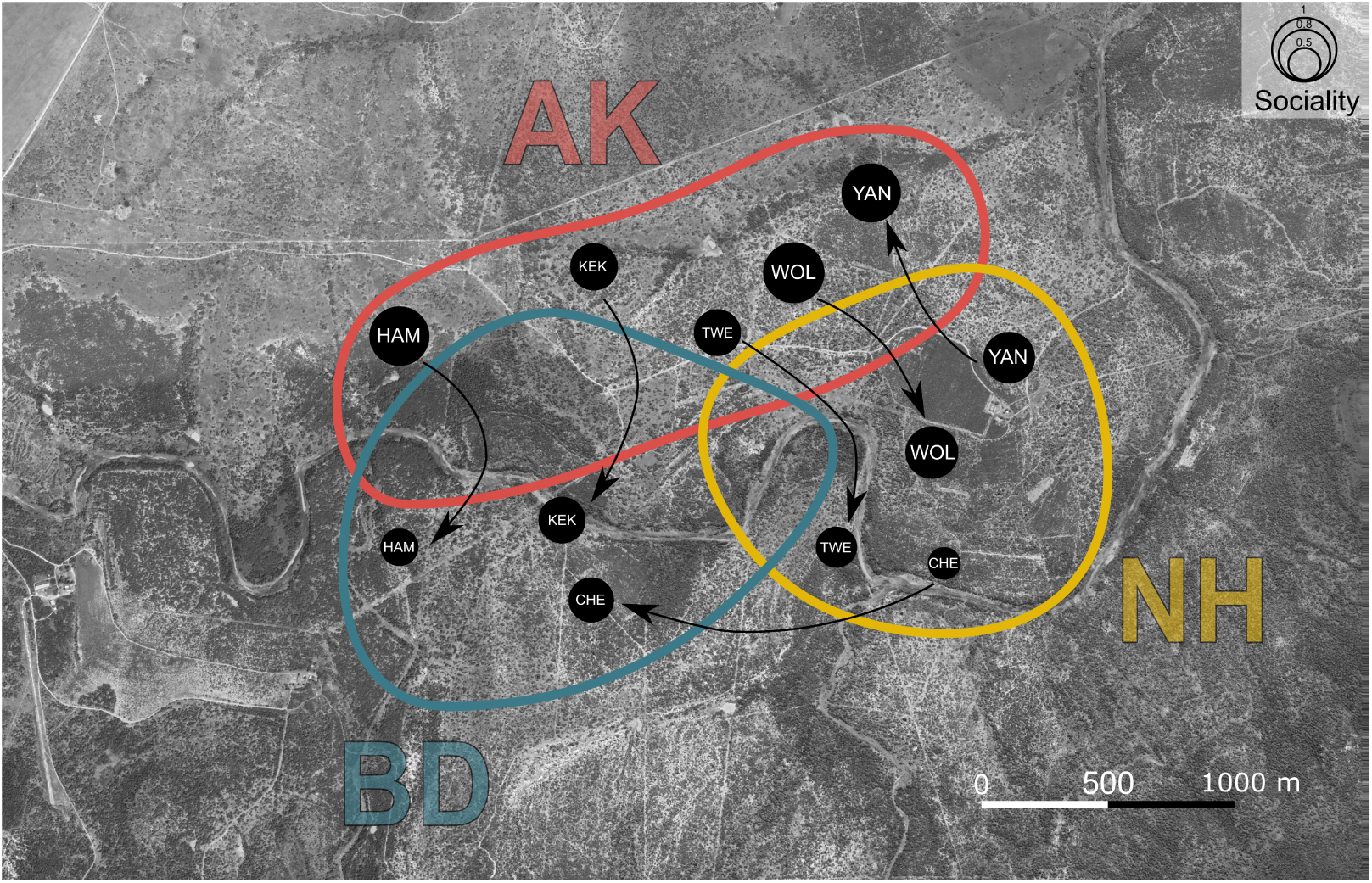
Dispersing map of males between groups. Map depicting the sociality of dispersing males between AK (red), BD (blue) and NH (yellow). Arrows represent the direction of dispersal. Each dot represents a male in a specific group. The bigger the dot, the more social the individual was. Sociality was computed from raw data. AK: Ankhase, BD: Baie Dankie, NH: Noha.

**Figure 3.**
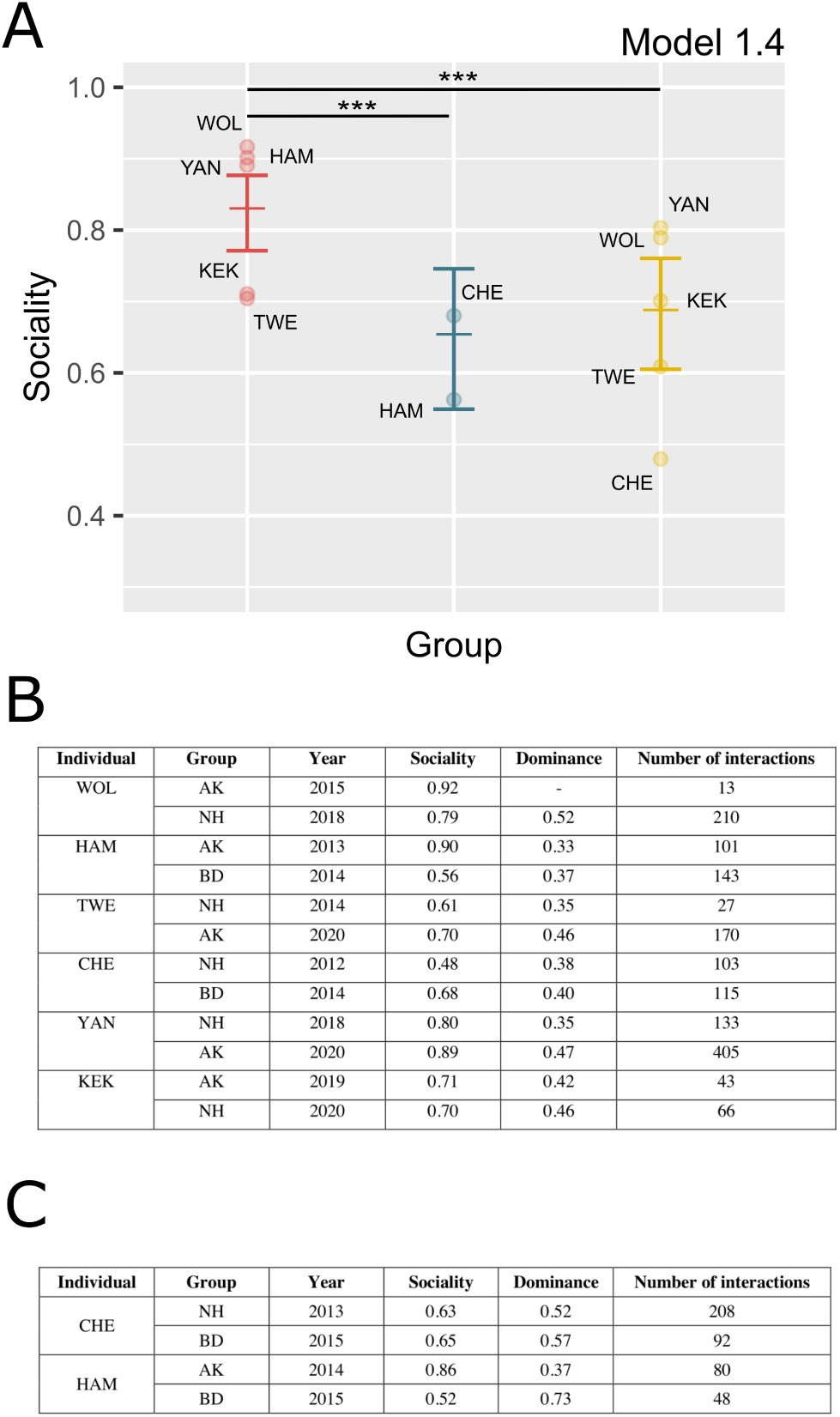
Adult males become less social when they disperse to BD and NH compared to AK. A) Mean sociality of dispersing males that dispersed to AK, BD and/or NH. Each dot represents individual sociality during its first year of arrival in the group, computed from raw data. The small horizontal bars denote the predicted mean of male sociality within their first year present in a group as predicted by Model 1.4. Error bars denote the standard error of the mean (SEM). (***) denotes the significant effects at the *α <* 0.001 level. B) Table displaying sociality indices of dispersing males during their first year of presence in each group. C) Table displaying sociality indices of dispersing males during their second year of presence in each group. Dominance was computed from Elo-rating method. AK: Ankhase, BD: Baie Dankie, NH: Noha

Model 1.4 (Table 1) highlighted, once again, an effect of group identity (*LRT_Group_* = 29.29, *p <* 0.001), with AK being the group where males were significantly more social after their dispersal (*AK > BD ∼ NH*: *AME_AK−BD_* = 0.18, *p <* 0.001; *AME_AK−NH_* = 0.15, *p <* 0.001; *BD − NH*: *p* = 1.00). According to Model 1.4 - for a specific year of our study – the sociality of one dispersing male arriving in AK would be 18% higher than if he had dispersed to BD, and 15% higher than if he had dispersed to NH (Figure 3A).

Regarding males who stayed a second year within each group and in agreement with our preceding results, we observed that HAM had a sociality 1.65 times higher in AK than in BD, while CHE had a similar sociality in BD and in NH (Figure 3C).

### Question II: The effect of group identity is confirmed within all socio-demographic categories

Overall, Model 1.5 (Table 1, Table S5) predicted that sex, age and hierarchical rank had a significant effect on individual sociality. Most importantly, these effects were similar within each group. Indeed, males sociality was significantly lower than females sociality (*AME_M−F_* = *−*0.09, *p <* 0.001), juveniles’ sociality was higher than adults’ sociality (*AME_Juv−Ad_* = 0.02, *p <* 0.001) and sociality significantly decreased as the hierarchical rank increased (*AME_hierarchical−rank_* = *−*0.07, *p <* 0.001) (Figure 4A, Figure 4B). The influence of *Sex* on sociality was about six times stronger than the one of *Age* and two times stronger than the effect of *Hierarchical rank*. These global effects were similar within each group as we found no significant effect of the interactions between *Group* and *Sex*, *Age* or *Hierarchical rank* (*Group*Sex* : *p* = 0.25; *Group*Age*; *p* = 0.55; *Group*hierarchical rank* : *p* = 0.62).

**Figure 4.**
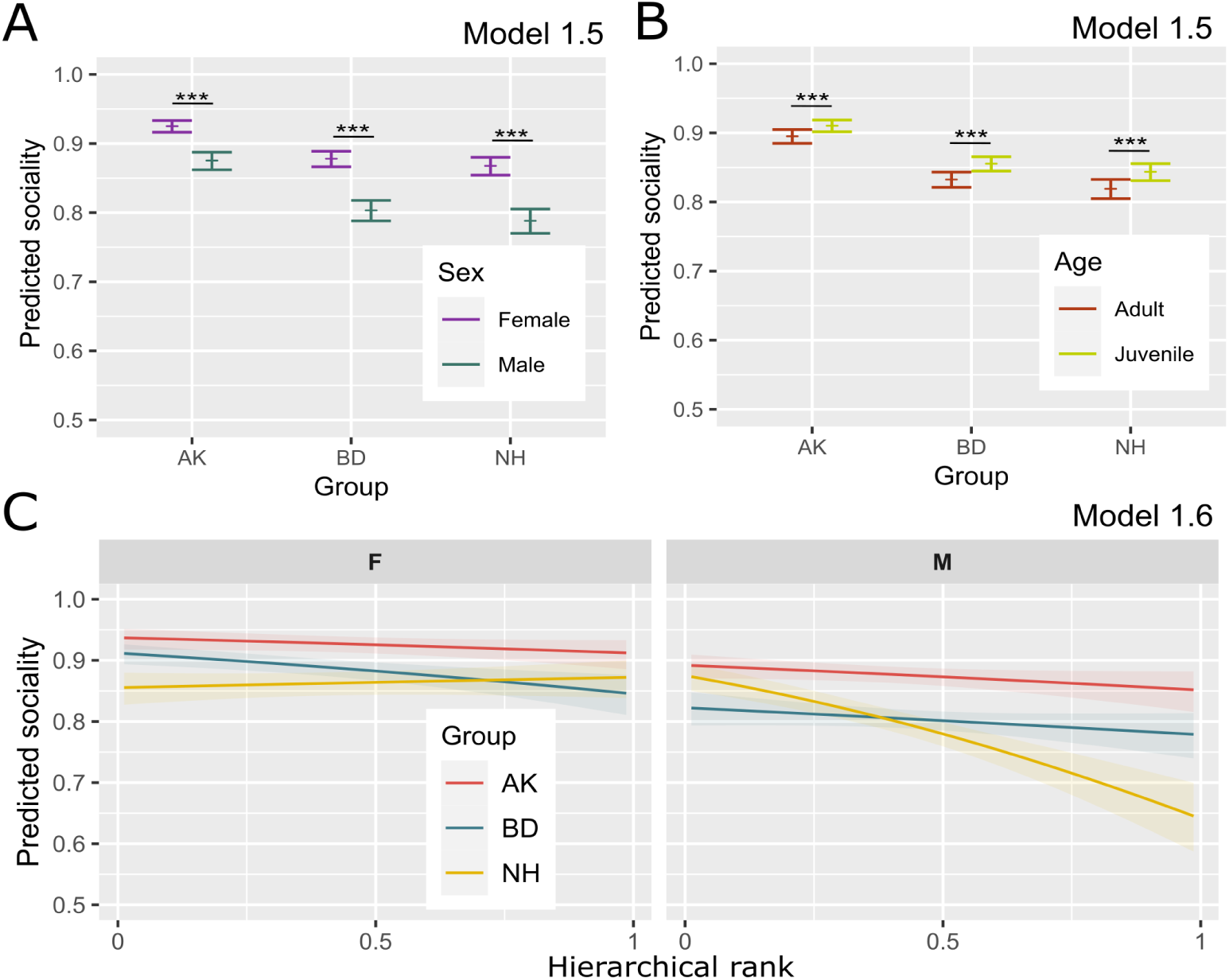
The effect of group identity is confirmed within all socio-demographic categories. A) Predicted sociality of individuals according to their sex in each group. The small horizontal bars denote the predicted mean sociality of males and females within each group according to Model 1.5. Error bars denote the standard error of the mean (SEM). (***) denotes the significant effects at the *α <* 0.001 level. B) Predicted sociality of individuals according to their age in each group. The small horizontal bars denote the predicted mean sociality of adults and juveniles within each group according to Model 1.5. Error bars denote the standard error of the mean (SEM). (***) denotes the significant effects at the *α <* 0.001 level. C) Differential effect of hierarchical rank in females’ (F, on the left) and in males’ (M, on the right) sociality within each group. Graphical representations are based on Model 1.6. Shaded areas denote 95% confidence interval (CI). AK: Ankhase, BD: Baie Dankie, NH: Noha.

More detailed analysis revealed a significant interaction of *Group*Sex* hierarchical rank* (*LRT* = 21.71, *p <* 0.001). The results of Model 1.6 (Table 1) showed that the way the hierarchical rank influenced the effect of sex on sociality was different in NH compared to AK and BD (*NH-AK*Female*hierarchical-rank* : *p* = 0.002; *NH-BD*Female*hierarchical-rank* : *p <* 0.001; *BD-AK*Female*hierarchical-rank* : *p* = 1.00). In Figure 4C, this result is illustrated by the parallel evolution of sociality according to hierarchical rank in both males and females in BD and AK. However, the curve was greatly modified in NH males. Indeed, in NH, an increase in males hierarchical rank induced a strong decrease of their sociality (*β_hierarchical−rank_* = *−*1.37, *p <* 0.001). We note that this interaction only influenced the relational order between BD and NH individuals’ sociality, but not the relationship between AK and NH, such that, NH sociality curve never crossed AK curve on Figure 4C. It is still worth mentioning that this interaction influenced the magnitude of the difference between NH and AK individuals’ sociality - for males of low dominance (close to 0), the difference in sociality between AK and NH was not significant (*p* = 1.00). Similarly for the highest rank females, no differences between NH and other groups could be detected (*AK − NH*: *p* = 0.60; *BD − NH* = 1.00). We additionally found a significant interaction between *Age*, *Sex* and *Hierarchical rank* (see Figure S1).

### Question II: The propensity to be affiliative is less directed towards related individuals in AK and BD

In Model 2.0 (Table 1), we found a significant effect of group identity on the matri-love index (*p <* 0.001) (Figure 5A). Matri-love index indicates the directionality of the sociality of an individual, directed either towards individuals from the same matriline (related individuals) or other individuals: the higher the value, the higher the propensity to be affiliative toward related individuals. NH individuals had a significantly higher matri-love index compared to individuals in AK and BD (*β_AK−NH_* = *−*0.07, *p <* 0.001; *β_BD−NH_* = *−*0.04*, p* = 0.013; *AK − BD*: *p* = 0.19). Model 2.1 (Table 1), revealed that the influence of hierarchical rank on matri-love was different across groups at a significance level of 0.05, however posthoc analysis failed to identify which group(s) differed from others (*Group ∗ hierarchical − rank*: *p* = 0.043; *NH − AK ∗ hierarchical − rank*: *p* = 0.86; *NH − BD ∗ hierarchical − rank*: *p* = 0.157; *AK − BD ∗ hierarchical − rank*: *p* = 1.00). Figure 5B illustrates the effect of hierarchical rank on individual matri-love within each group and suggests that the interaction may emerge from a stronger effect of hierarchical rank on NH individuals’ matri-love.

**Figure 5.**
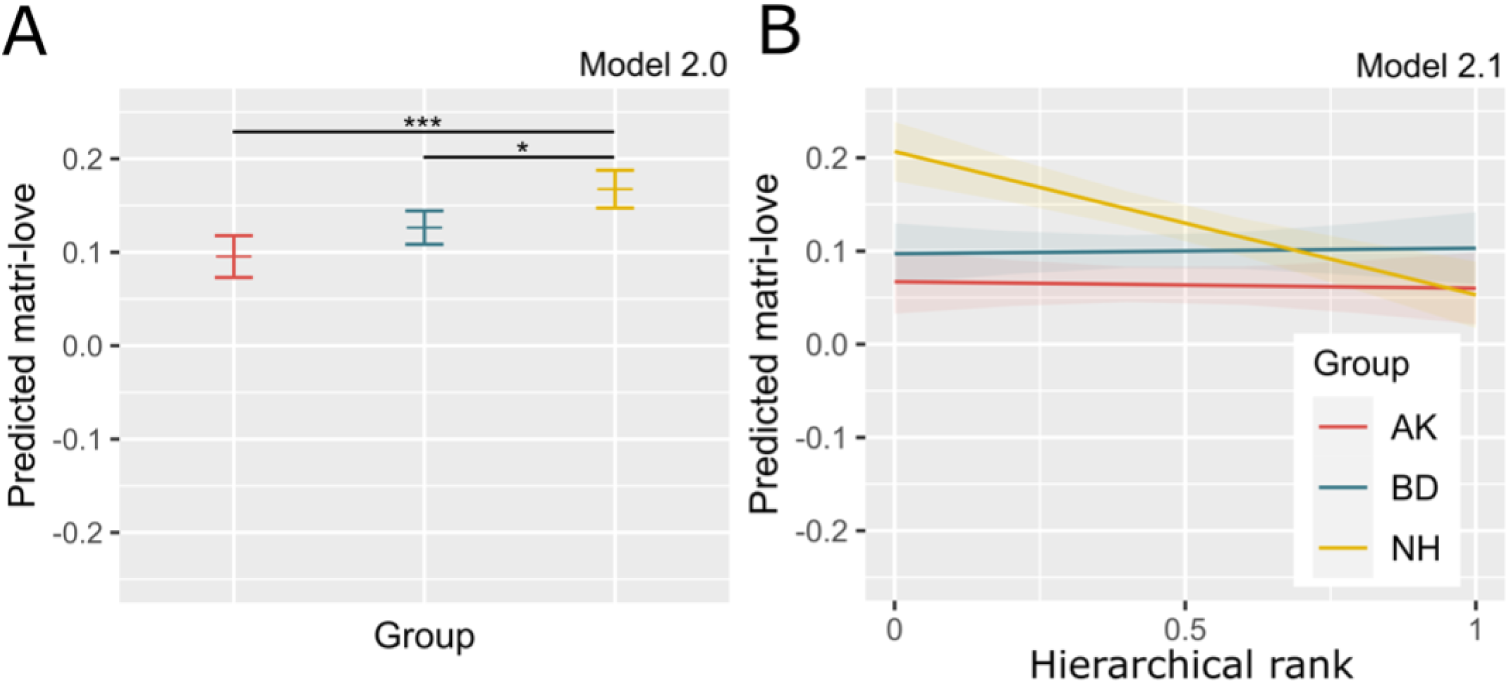
The propensity to be affiliative is less directed towards related individuals in AK and BD. A) Predicted matri-love per group. The small horizontal bars denote the predicted mean of matri-love as computed from Model 2.0. Error bars denote the standard error of the mean (SEM). (***) denotes the significant effects at the *α <* 0.001 level. (*) denotes the significant effects at the *α <* 0.05 level. B) Differential effect of hierarchical rank on matri-love within each group. Graphical representations are based on Model 2.1. Shaded areas denote 95% confidence interval (CI). AK: Ankhase, BD: Baie Dankie, NH: Noha

### Question III: Grooming is more evenly distributed among individuals in AK

In agreement with the results of Model 1.0 regarding sociality index, we found in Model 3.0 (Table 1) that the degree of grooming reciprocity significantly differed across groups. Over the nine years of our analysis, according to a simple analysis (Model 3.0), grooming was more evenly distributed between individuals in AK rather than in BD and NH (*AME_NH−AK_* = *−*0.06, *p <* 0.001; *AME_BD−AK_* = *−*0.07, *p <* 0.001). Grooming was also significantly more reciprocal between NH rather than BD individuals (*AME_BD−NH_* = *−*0.01, *p <* 0.001) (Figure 6A). Similarly to our observations for sociality, the results of Model 3.1 (Table 1) indicated that, in similar demographic environments, groups of vervets still significantly differed from each other (*LRT* = 150.459, *p <* 0.001) with the same results as in Model 3.0 (*AK > NH > BD*: *AME_NH−AK_* = *−*0.05, *p <* 0.001; *AME_BD−AK_* = *−*0.07, *p <* 0.001; *AME_BD−NH_* = *−*0.02, *p <* 0.001) (Figure 6B). Overall, as the relative number of males or adults increased in the group, the grooming reciprocity decreased (*AME_Sex−ratio_* = *−*0.03, *p <* 0.001; *AME_Age−ratio_* = *−*0.03, *p <* 0.001). At a more precise level, we found that the interaction between Sex-ratio*Age-ratio varied across groups with an opposite pattern between AK and the two other groups (*LRT_Group∗Sex−ratio∗Age−ratio_* = 13.874, *p <* 0.001; see Figure S2 for details).

**Figure 6.**
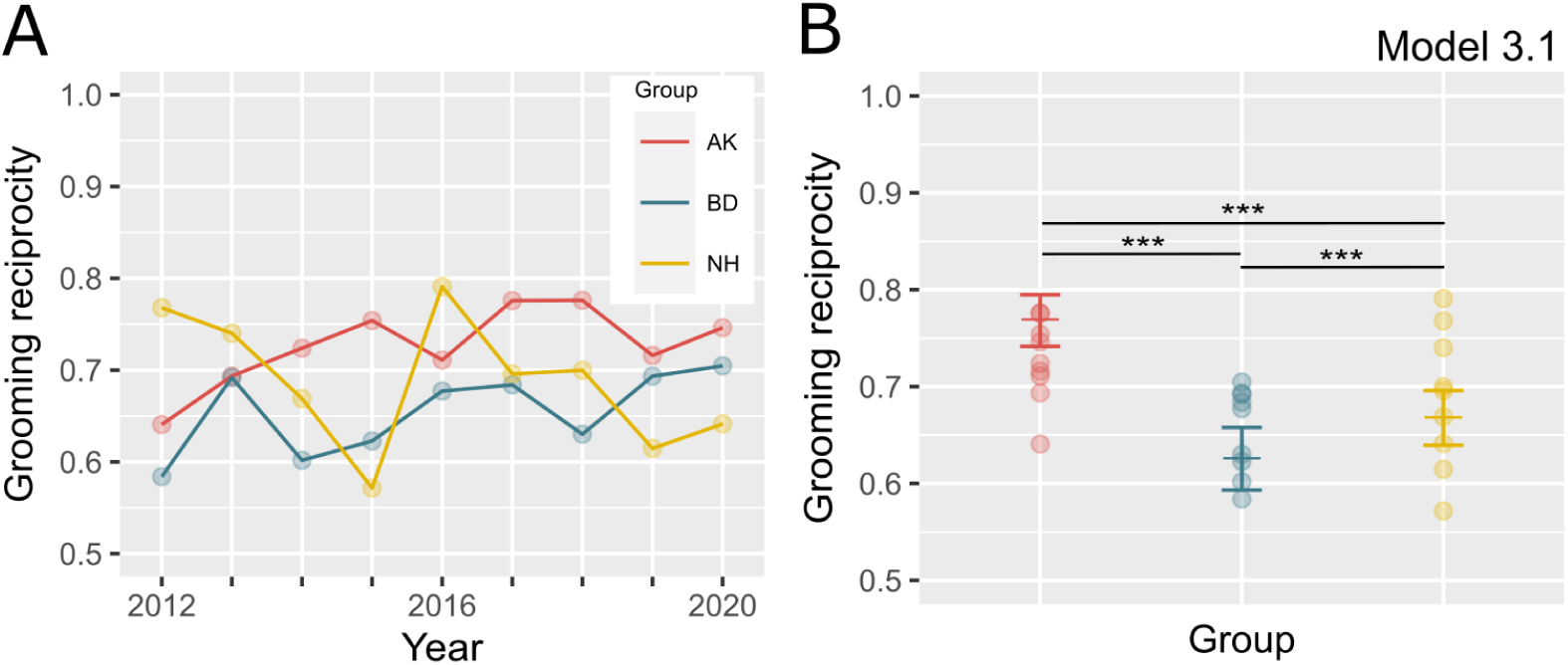
Grooming is more evenly distributed among individuals in AK. A) Dynamics of the level of grooming reciprocity in AK, BD and NH across years. Dots represent the grooming reciprocity of each group for each year of the study as computed from raw data. B) Predicted grooming reciprocity in AK, BD and NH for similar demographic environments. The small horizontal bars denote the predicted mean of grooming reciprocity within each group as predicted by the Model 3.1. Error bars denote standard error of the mean (SEM). Dots represent the grooming reciprocity of each group for every year of the study as computed from raw data. (***) denotes significant effects at the *α <* 0.001 level. AK: Ankhase, BD: Baie Dankie, NH: Noha

## DISCUSSION

Despite the large body of research on primate culture^12,1,53^, there is still a lack of evidence regarding the existence and stability of social traditions, especially in the wild^54,31,37^. Our research contributes filling this gap by investigating inter-group differences in sociality and their stability over nine years of data collected on three neighbouring groups of wild vervet monkeys. We found that individuals from one group – AK – were globally more affiliative than individuals in the two other groups, BD and NH. Taken together, our results show that these stable group differences in social dynamics are independent from demographic variables and are unlikely to be explained by genetic and ecological differences alone. Moreover, we report that males who dispersed between our study groups adjusted their sociality, becoming more social when they integrated into AK compared to the other groups and conversely becoming less social when they joined BD or NH, suggesting social conformity (as defined in^55^). Combined, these results suggest that these lasting group differences in basic social behaviours can be considered as social traditions in wild vervet monkeys.

### Question I: sociality is predicted by group identity

#### AK individuals are more social

In agreement with our first hypothesis (I), we find that individuals in one group - AK - had a higher propensity to be affiliative than individuals in the two other groups and that this effect was stable over the nine years of study despite the yearly fluctuations of each group’s sociality. These differences were not due to trivial differences in demography, as group differences were maintained for similar group size, age, and sex ratios in our modelling. As an example, a change in one standard deviation value of sex or age ratio impacted group sociality ten times less than group identity did.

#### Origin of group sociality differences: demographics, ecology and genetics sources of behavioural variability

The list of demographic variables influencing animal social behaviours is non-exhaustive^56^ and we do no pretend to have captured the influence of the wide scope of potential demographic factors. For example, due to our data structure, we were unable to control for the number of births and deaths, which are known to influence social behaviour in vervet monkeys^57^ and in gorillas^58^. Even if we cannot completely exclude that one part of group variability in sociality might be due to variations in demography, the temporal scope of our study combined with the variability in each group demography enable us to exclude demography as the main and only source of the observed differences in sociality.

Similarly, as argued before^59^, it is impossible to completely dismiss the potential effect of local ecological variability on sociality. Thus, even if ecological variability is unlikely to be the main driver of sociality differences as our study groups shared a similar global habitat, we cannot exclude that groups’ territories may be heterogeneous in resource availability and quality^60^. Variability in the ecological structure of each territory combined with the presence of specific predators may also have formed a heterogeneous “landscape of fear” (LOF)^61,62^ where “prey perceive and respond to predation risk across both space and time simultaneously”^63^ differently in each territory. Vervets are able to specifically recognize different predators^64^ and their perception of predation risk (mainly from baboons and leopards) has been shown to influence their use of space^65^. We can then hypothesize that local differences in the LOF may have partially influenced the observed group differences in social behaviours.

Behavioural variability in primates can also be linked, to some extent, to genetic variability itself^66^. For example, Langergraber et al.^67^ highlighted how behavioural dissimilarities reported by Whiten et al.^12^ between wild groups or populations of chimpanzees were, for the majority (52%), significantly correlated with genetic dissimilarity between these groups although no causality was here advanced. For the only social type of tradition reported in this paper, namely hand-clasp grooming, a genetic origin could not be explicitly excluded. At the neurobiological level, neurogenetics studies have highlighted how polymorphisms of coding or regulatory regions can lead to inter-individual variability in aggressive behaviours in primates, for example through the dopamine D4 receptor and the serotonin transporter^68^. However, in another study on wild orangutans^69^, the proportion of geographic variations in traditions explained by genetic or environmental differences among populations was very low. A specific quantification of genetic dissimilarities between our groups of vervets could thus help future IVP-based studies to potentially exclude the hypothesis that genetic variability is partially linked to the observed behavioural variability. However, as highlighted by Schuppli et al.^59^, “in practice, genetic components do not rule out the presence of social learning”, so that even if we are unable to completely disregard the potential influence of genetics in our study it does not exclude the eventuality that social learning has a significant role in the observed sociality differences between our groups of vervets.

#### Transmission through social learning to dispersing males: a case of social conformity?

We reported that dispersing males were always more social in AK rather than in BD or NH. This effect was independent of the year of dispersal and of individual hierarchical ranks. These results were obtained from a limited set of individuals for which we had observation for a sufficient amount of time in the different groups. Nonetheless, these findings suggest that the previously mentioned differences in sociality between groups might be socially maintained.

Conformity can be defined as “how individuals alter their behaviour to be similar to that of others”^70^ and is central in recent reflection on animal behavioural variability^70^ and more specifically animal^71,72^ and human traditions^73^. Our result is consistent with the previously suggested conformity effect in dispersing males in wild vervet monkeys^47^. In the van de Waal et al. study^47^, dispersing males adapted their foraging behaviour to the local norm rather than following their natal preference. This type of conformity relates to “Aschian conformity”^74^ adopted through “overriding of personal knowledge or behavioural dispositions by countervailing options observed in others”^75^. Later, such conformity was also described in the food preference of female vervet monkeys^55^ and was reported as a potential “social conformity” where “individuals act like others not to achieve an informational function, but instead to achieve a social function that derives from simply “being like others’”” as previously suggested by de Waal’s Bonding and Identification-based Observational Learning theory^76^. We can hypothesize that a “chameleon effect”^77^ might be at stake with males mimicking others in their social behaviour. This behavioural mimicry would act as a social glue and allow males to get better integrated in their new group. Such social conformity could be a potential vector of intra-group integration^55^. In the perspective of our study, social conformity could then play a similar role in the observed changes of adult males’ sociality. Indeed, intra-group integration is a major challenge to dispersing males, who must join another group to mate and survive. Even after dispersal, males may inherit the social bonds of certain conspecifics serving as role models^78^. Further experiments should be designed to assess if and how social dynamics can be socially transmitted and if it could assist in the intra-group integration of dispersing males. Interestingly, cases where dispersing males bring their individual knowledge to their new group after dispersal have also been reported, however, such observation often concerns food-related and material traditions^79,80,81,82,83^ and have rarely been reported in social traditions^54,84,83^.

Overall, our results add to previously published studies which offer evidence for the existence of social traditions in bonobos ^85^, chimpanzees^86,87,88^, capuchins^54,89^, spider monkeys^32,31^ and baboons^16,37^.

#### A female-driven effect?

Interestingly, our results suggested that group differences in sociality may be driven by females. Indeed, differences in sociality between groups were stronger when groups contained a female-biased sex ratio. Regarding the literature, two main hypotheses (i; ii) emerge. The first hypothesis (i) suggests that AK females might be more genetically linked to each other than females in BD and NH. According to Hamilton’s theories on the “selfish gene”^90^, increasing the number of females in a group, by highly increasing the over-all genetic link between individuals, would therefore favour high levels of inter-individual affiliative behaviours. This genetic hypothesis is however contradicted by the decrease of sociality in BD and NH sociality when the groups had female-skewed sex ratios, and is thus insufficient to entirely explain our results. The second hypothesis (ii) proposes that increasing the relative number of females in a group might strengthen behavioural group differences, with females acting as “models” for social learning. Adult females (here the philopatric sex) form the social core of the groups ^45^ and may act as preferential “models” for social learning in wild vervet monkeys^46,48^ (but see^49,50^). Moreover, previous studies highlighted the females’ role in the spread of social traditions to incoming adult males^37^ and it has been shown that maternal affiliative relationships may be socially transmitted to offspring, similarly to the determination of offspring rank by maternal rank in macaques and in hyenas^91,92^. More precisely, longitudinal observational studies and cross-fostering experiments reported similarities in behaviour such as rate of infant rejection and social contacts between mothers and daughters in vervet monkeys^27^ and rhesus macaques,^28,29^ suggesting that these similarities may be the result of daughters’ early experience. Regarding the maintenance of group differences in sociality, Ilany & Aķcay^93^ proposed a model of social inheritance from parents. According to their model, juveniles are likely to form social bonds with their parent’s conspecifics with whom they had positively interacted before. This model has been validated in elephants^94^ but not yet in vervet monkeys^95^.

#### Effect of habituation on sociality

We observed a constant increase in sociality across the nine years of the study for each group. This result could be due to the gradual habituation of vervets to human presence, where habituation is defined as “the process by which repeated exposure to humans results in a gradual reduction in animals’ fearful response, until animals no longer perceive humans as a direct threat”^96^. At the start of our study in 2012, the three groups were less habituated to observers so human presence might have stressed monkeys, impacted their behaviour and generated more conflicts. Data collection was also more difficult compared to nine years later where monkeys were highly habituated to human presence. It is thus possible that loud conflicts were easier to spot than silent grooming at that period, and especially if the monkeys were hiding from researchers when doing it. Another hypothesis would be that growing interactions with humans resulted in enhanced sociality in monkeys as enculturation in great apes improves their socio-physical cognitive skills^97,98^**^?^**. Another potential explanation would be the existence of two stable states of the overall sociality: one before 2015 and one after 2015. Interestingly, 2015 was a year of unusual drought (see Newcastle station^99^), and such environmental perturbation may have elicited changes in groups social dynamics^100^.

### Question II: Group differences in sociality are not explained by simple effects of individuals’ socio-demography or kinship

#### An even increase of sociality across all socio-demographic classes in AK

Our results indicated that AK high sociality was not due to a specific socio-demographic category but rather emerged from a global increase of sociality within every socio-demographic category. Indeed, sex, age and hierarchical rank had similar effects on sociality in the three groups.

#### The effect of sex, age, and hierarchical rank on sociality

Our analysis confirmed the robustness of previous findings regarding socio-demographic influences on sociality and report unexpected effects. First, we found that females were more social than males. This result corroborates previous studies showing that females tend to be more involved than males in grooming and play interactions and less involved in agonistic interactions^23,101,22^. Additionally, we found that juveniles were generally more social than adults. This could be due to the fact that juveniles are more often involved in play interactions than adults, but equally involved as adults regarding grooming interactions or agonistic interactions ^23^. However, the effect of age on sociality was rather weak and may thus have a limited role in intra-group variation. Nevertheless, we found a negative effect of hierarchical rank on sociality. This result suggests that, even if high-rankers have been shown to be more involved in both grooming and agonistic interactions (acting more as aggressor and receiving more grooming than low-rankers)^23^, the relative increase in aggression might be greater than the relative increase in grooming interactions. In particular, we found that this effect was even stronger in NH males compared to the other groups.

#### AK individuals are not more affiliative towards related ones

Our study also suggested that AK high sociality did not emerge from biased social interactions towards related individuals. However, in contrast with our hypothesis, we found that NH individuals had a higher tendency to be affiliative towards related conspecifics rather than unrelated ones, compared to AK. This group difference may derive from the behaviours of low-rankers in NH. However, as our knowledge of matriline cannot extend prior to 2010, we cannot exclude the fact that what we defined as “unrelated” conspecifics may be relatively more related in our group than in another. For example, two adult females identified as unrelated in 2010 could actually be sisters so that we underestimate how closely related individuals from these matrilines are. Combined with the effect of the hierarchical rank on NH males’ sociality, this result suggests that the hierarchy had a greater impact on social behaviours in NH compared to AK and BD.

#### Influence of “keystone” individuals?

In the present study, some individuals were present from the start to the end of the study within a single group, among them one individual in each group (GUGU for AK, OULI for NH, UPSS for BD) had a mean elo-rating higher than 0.8 over the whole study. It is possible that these individuals had a significant impact on the sociality of the group, via a substantial influence on other members of the group (“keystone” individual^102^). We argue that the field would greatly benefit from the precise dissection of the influence of individuals’ personalities on group-specific primate social behaviours. Indeed, several studies detected the presence of “key” individuals to have a significant impact on group-specific “collective behaviours”^102,103,104,105,106^. Their impact has been especially studied in the enhancement of cooperative behaviours^106^.

### Question III: Grooming is more evenly exchanged among AK individuals

Interestingly, our results on grooming reciprocity corroborated the ones on sociality, both highlighted the specificity of AK global social dynamics. We found that grooming was exchanged more evenly between AK individuals rather than between BD or NH individuals. This difference between groups was however less stable over time than the difference we found in sociality. Grooming has been postulated as an exchange currency according to the Biological Market Theory (BMT) (Baboons^107^, Chimpanzees^108^, Rhesus macaques^109^, Bonobos^110^, Tibetan macaques^111^, Vervet monkeys^52^). According to this theory, grooming (i.e. good) is exchanged with another commodity (i.e. service) based on the law of supply and demand and predicts differences in rank and high overall competition to influence negatively reciprocity of grooming^112^. Within the BMT, this result could be interpreted as a less tense state of the biological market in AK than in BD and NH, where high-rankers have a lower power over low-rankers (i.e. less dominance-based disproportion in resource access)^107^. Our measure of grooming reciprocity is however unconventional, as we were unable to obtain consistent data on the more conventional direct reciprocity - number of grooming bouts within a single, time-condensed, grooming interaction. Further studies using direct reciprocity and considering the time span of each grooming bout^113,114,110^ should be considered to confirm these results.

In conclusion, our study reports the existence and stability of different degrees of sociality and grooming reciprocity in three groups of wild vervet monkeys. Such differences in social dynamics could not be explained by differences in group demographics, and they impacted every socio-demographic class. These differences in sociality across neighbouring groups of wild vervet monkeys cannot be explained by genetic and environmental differences alone. Thus, part of this variability may emerge from social learning, emphasized by the fact that dispersing males conformed to the local sociality of the group they joined. Thus, these results provide new insights on the existence of social traditions in wild monkeys. In accordance with Schuppli & van Schaik^59^, we believe that researchers have only discovered the tip of the primate tradition iceberg so far, mainly composed of complex technical behaviours, while missing information about the base of this iceberg. With this study, we contribute to filling this gap, suggesting that this base is composed of basic skills from the social realm.

### Limitations of the study

In our study, we used data collected *ad libitum*, a method for which it is difficult to accurately assess the sampling effort (but see Canteloup et al.^115^ for high correlation of social networks assessed via *ad libitum* and focal animal sampling). One limitation of *ad libitum* sampling is that shier individuals may be sampled less often than others, and thus appear less social than they are, which could lead to a potential bias in social networks. Additionally, although our study groups share the same environment with genetic flux between groups, we do not have precise data on ecological and genetic variability. Future work on the ecological and genetic variation of our study groups would bring some elements regarding potential fine-grained differences (e.g. “landscape of fear”) in our study groups’ territories. Finally, the influence of keystone individuals on group social dynamics is still poorly known and more work on animal personalities are needed to disentangle between such effects.

## ACKNOWLEDGMENTS

We thank the onsite managers, Albert Driescher and Arend van Blerk and all the field assistants, Master students, PhD students and postdocs who collected the data over the study period. We are grateful to the van der Walt family for giving us the permission to conduct the study on their land. We thank Pooja Dongre for her advice on the Elo-rating calculation and Rachel Harrison, Fédéric Schütz, Christof Neumann and Cristian Pasquaretta for their statistical advice as well as Leslie Ng for his help on English proofreading. This project was funded by the Swiss National Science Foundation (*P* 300*P* 3 151187, 31003*A* 159587, *PP* 00*P* 3 170624 and *PP* 00*P* 3 198913) along with Branco Weiss Fellowship–Society in Science and the grant ‘ProFemmes’ of the Faculty of Biology and Medicine, University of Lausanne (granted to EvdW) and by a Fyssen grant (granted to CC).

## CONTRIBUTION

Conceptualisation: Elena Kerjean, Erica van de Waal, Charlotte Canteloup

Data curation: Elena Kerjean Formal analysis: Elena Kerjean

Funding acquisition: Erica van de Waal, Charlotte Canteloup

Investigation: Elena Kerjean, Charlotte Canteloup

Methodology: Elena Kerjean, Charlotte Canteloup

Project administration: Charlotte Canteloup, Erica van de Waal

Resources: Erica van de Waal

Software: Elena Kerjean

Supervision: Charlotte Canteloup

Validation: Charlotte Canteloup

Visualization: Elena Kerjean

Writing – Original draft preparation: Elena Kerjean, Charlotte Canteloup

Writing – review & Editing: Elena Kerjean, Charlotte Canteloup, Erica van de Waal

## METHODS

### RESOURCE AVAILABILITY

#### Lead contact

Further information and requests for resources should be directed to and will be fulfilled by the Lead Contact, Elena Kerjean.

#### Materials availability

This study did not generate newly generated materials.

#### Data and code availability

Behavioural data and all codes use for statistical analysis have been deposited at zenodo.7997943 and are publicly available as of the date of publication. Accession numbers are listed in the key resources table. Any additional information required to reanalyze the data reported in this paper is available from the lead contact upon request.

## EXPERIMENTAL MODEL AND STUDY PARTICIPANT DETAILS

Data were collected over nine years (2012-2020) within the IVP, at Mawana game reserve (28°00.327S, 031°12.348E) in KwaZulu Natal, South Africa. We used the data collected in three neighbouring groups of wild vervet monkeys: AK, BD, and NH. These groups have been studied as part of IVP since 2010 and have been habituated to human presence since that time. We only included individuals that stayed in their group for at least six months during a given year in data analysis to ensure we had sufficient data per individual to analyse. Unless mentioned, we included individuals of both sexes and all ages in our analysis. Data collection was approved by the relevant local authority, Ezemvelo KZN Wildlife, South Africa.

## METHOD DETAILS

### Behavioural sampling and coding

We used data collected by trained research assistants and researchers using *ad libitum* sampling^116,117^. In such a case, no constraints are placed on the individuals for which data are recorded nor the period when the data are recorded. An interaction is defined here as one or more behaviours performed between two individuals (hereafter, dyad). On the basis of established ethograms of social behaviours (Table S1, Table S2, Table S3), we defined affiliative interactions as interactions including only affiliative and neutral behaviours and agonistic interactions as interactions including only agonistic and neutral behaviours. Our dataset included a total of 68 443 affiliative interactions and 16 259 agonistic interactions. In 2017, three months (June, July, August) of missing time of observation and/or affiliative interactions were excluded from our analysis. Observers recorded behaviours as point events. To assess sampling effort, we took the total amount of time spent in the field during field days as a proxy for the effort of *ad libitum* sampling in each group. We assessed this effort by computing the duration of observation of each group per day which we multiplied by an attenuating factor of the number of observers for each field session according to a binary logarithmic function, so that the weight of each observer decreased as the number of total observers increased (AK mean *±* SD: 2706 *±* 803, BD mean *±* SD: 3360 *±* 614, NH mean *±* SD: 3439 *±* 974). Before data collection, each observer passed an inter-observer reliability test (80 % agreement required between two observers).

### Demographics and socio-demographics

Group size was computed as the number of individuals included in our analysis for each group in each year (AK mean: 27 individuals *±* 8.5, range: [18-41]; BD mean: 50 individuals *±* 6, range: [43-62]; NH mean: 39 individuals *±* 7, range: [31-50]). The sex ratio was computed as the number of males divided by the number of females in each group for each year, so that the more the sex ratio increased, the greater the relative number of males was in the group. The age ratio was defined as the number of adults divided by the number of juveniles in the group, similarly, the more the age ratio increased, the greater the relative number of adults was in the group (Table S4). We classified individuals according to four age classes: newborn (from their birthday to the end of their birth year), infant (from the start of birthyear +1 until first birthday), juvenile (male: 1-5 years old; female: 1-4 years old) and adult. Regardless of their age, as soon as they gave birth (for females) or dispersed from their birth group (for males), individuals were considered adults. After the end point of the juvenile period (five years old for males, four years old for females) even if they had not met these criteria of adulthood, individuals were considered as adults. Hereafter, when individual level analyses were performed, newborns and infants were excluded due to the lack of a hierarchical rank for these individuals. We computed the hierarchical rank of all individuals using the elo-rating method^118^. To do so, we based our analysis on the Elorating 0.46.11 package of R^119^. The specificity of this method lies in the fact that it updates individual hierarchical rank after each conflict: winners increase their rating while losers decrease their rating. The magnitude by which the rating changes depends on the expected outcome of the conflict (i.e., the difference between the previous ratings of the two opponents). We specified the relationship between the rating difference and the winning probability as a sigmoidal curve, following^120^. The k parameter of the maximal amplitude of change after an interaction has been set to 50. The initial rating for all individuals was 1000.

### Social indices

#### Sociality indices

The sociality indices are here defined as the proportion of affiliative interactions (i.e., the number of affiliative interactions divided by the sum of the number of agonistic and affiliative interactions). We computed two sociality indices: a group sociality index and an individual sociality index. The group sociality index was computed as the proportion of affiliative interactions in a specific group during a year, regardless of the individuals involved in each interaction. The individual sociality index was computed as the proportion of affiliative interactions per individual. Both sociality indices ranged from 0 to 1. A sociality index close to 0 means that individuals were more agonistic than affiliative, and conversely when close to 1, individuals were more affiliative than agonistic. Affiliative behaviours mainly included grooming events (80% of all affiliative behaviours), so each interaction was treated as qualitatively equivalent. To take into account the level of aggression of each agonistic interaction, we attenuated the weight of mild agonistic interactions (without direct contact) by a factor of 0.5. Sever agonistic interactions were classified as: attack, bite, chase, fight, grab or hit (Table S2).

#### Matri-love index

We computed a matri-love index at the individual level. The matri-love index quantified the directionality of individual sociality: either toward individuals from the same matriline or not. A matriline, includes mothers and their offsprings. All siblings are from the same matriline. Each matriline starts from one of the adult females present at the starts of IVP.

We computed two sociality scores for each individual: the first one included only the interactions involving individuals from the same matriline while the second one included only the interactions between individuals from a different matriline. We defined the difference between these two measures as the matri-love index. The matri-love index is lower than 0 when sociality is higher with individuals from a different matriline and higher than 0 when sociality is higher with individuals from the same matriline.

#### Grooming reciprocity index

We computed one grooming reciprocity index at the group level. In total, we included 63 325 grooming behaviours in our analysis. To compute the grooming reciprocity index, we built a matrix of reciprocity (M) between individuals of a single group during a specific year. The reciprocity index (R) was then calculated with the following formula:

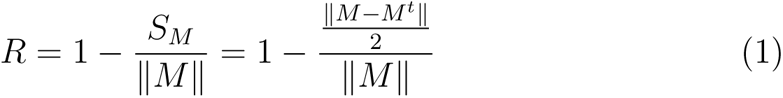

where ∥ *M* ∥ *≠* 0 and with *S_M_* the distance of the matrix M to the sub-vectorial space of symmetrical matrices and *M^t^*denotes the transposed matrix of M with ∥ ∥, being the euclidean norm. R is thus a measure of the similarity between the upper triangular part of the matrix M and its lower triangular part.

## QUANTIFICATION AND STATISTICAL ANALYSIS

Statistical models (Table 1) were assigned a number in the form x.y, where x defines the dependant variable used in the model - either sociality (1), matri-love (2) or grooming reciprocity (3) - and y defined the sequential order of analysis. All models, unless mentioned, used a binomial distribution of the error terms with a logit link function. This structure defines our dependant variables by a logistic function (i.e., inverse of logit function). To help model convergence and to make interpretation of estimates easier, continuous covariates were Z-transformed to a mean of 0 and a standard deviation of 1 prior to model selection^121^. One exception was made for the variable describing the Elo-rating as it is bounded between 0 and 1. Final models were compared to a corresponding null model. To assess whether the inclusion of predictors improved our model or not we built null models including only controls and random effects. We used the function *anova* with the test parameter set to “LRT” to compare two GLMMs (generalized linear mixed models). To assess the relevance of random effects we used the lowest Akaike information criterion (AIC) to we compare GLM and GLMMs (with a difference higher than 2^122^). We used the function *ranef* to assess the effect of random slopes. We chose random effects with at least five levels^123^.

### Model construction

We based our model selection on a forward selection strategy using the AIC as a selection criterion^124^. We first interpreted the value of estimates of the main effects before including significant interactions in the models. We interpreted only significant interactions which included the fixed effect Group, as our interest was focused on the effect of group identity. To assess the significance level of each of the predictors, we compared the full model to a model with one omitted predictor via a likelihood ratio test (we used the function *drop1* with the argument test set to “Chisq”). Post-hoc analysis on categorical variables was then performed using the Bonferroni p-value adjustment^125^. After a first interpretation, we tested interactions using *drop1* function and performed post-hoc analysis on these interactions. In models including significant interactions, we re-evaluated the effects of the main effects that were not included in the interactions, and only mentioned changes if the estimate changed from more than 2% or if the significance level changed. Then, we interpreted the main effects of the predictors part of the interaction as marginal effects conditional on the value of the second predictor being equal to its intercept value.

### Model evaluation

For each final model selected, we performed a two-stage evaluation. First, we verified multi-collinearity, using the function *vif* from the package *car* 3.0.12, measuring the variance inflation factor (VIF) of each fixed effect in our model. We made sure that generalized VIF (GVIF) values of each of the continuous variables and the square of *GV IF* ^1^*^/^*^2d^*^f^* for each categorical variable were lower than 5^126^. Secondly, we visually checked the distribution of the residual deviance according to the predicted values of our dependant variable^127^. We made sure this distribution did not show any specific pattern.

### Effect of predictors in GLMs and GLMMs

In the specific case of non-linear regression, GLMs and GLMMs, we obtained varying effect of predictors across the range of these predictors. The direction of the effect of the predictor was induced from the log-odds. To measure for the intensity of each effect we discussed the average marginal effects (AMEs) of each predictor and/or the estimated marginal means (EMMs) of the response variable for different values of each predictor. AMEs were obtained from the function *margins* from the package *margins* 0.3.26^128^, while EMMs were obtained with the function *ggemmeans* from the package *ggeffects* 1.1.2^129^. When results from statistical analysis are mentioned for categorical variables, in such a way: A-B, the second category (i.e., B) refers to the category defined as intercept in our analysis. We ran all analyses in R Studio version 1.3.1093 using R version 4.1.2^130^. Statistical parameters for each models are reported in Table 1.

### Question I

In order to have a global understanding of the differences in sociality between groups across years, we first computed a simple model which isolated *Group* as as the only main predictor of group sociality (Model 1.0). In this model, we used *Year* as a random intercept to take into account latent variables we did not collected such as temperature or food availability at each specific year and to assess the stability of group differences. In order to assess the effect of time in each group, and because we cannot expect *Year* to have a linear effect, we then constructed a model using *Year* as a random slope, keeping *Group* as a fixed effect (Model 1.1) and compared it to a model without *Year* as a random slope.

Next, we constructed a more complex model to control for the group demographic (sex ratio, age ratio and group size) and experimental (time of observation) variables, using *Year* as a random effect (Model 1.2). Due to a high collinearity of *Group Size* (20 *> V IF >* 10), and as the correlation between the group size and other variables in our model was low to mild (*Group* : 0.45, *Sex ratio* : 0.15, *Adult ratio* : 0.42), we kept this variable as a control in our models. To have a more precise understanding of the socio-demographic conditions under which group differences in sociality was modified we then included significant interactions (Model 1.3).

Because some males dispersed between our study groups over the course of the nine years study, we could investigate whether their sociality changed between the group they quit and the group they integrated. We included in the analysis (Model 1.4) behavioural data collected of six males who dispersed in between our three study groups and for which we had sufficient data. We calculated the mean group sociality as the mean of the sociality of the three groups during a year. As the mean group sociality during the arrival year of the males could be globally influenced by yearly ecological changes, we included it as a control variable in the model. As this analysis was performed at the individual level, individual identity was included as a random intercept. We removed individual hierarchical rank from the model as its effect on individual sociality was not significant (Log-Odds = *−*3.87, Confidence Interval : [*−*8.37, 0.63], *p* = 0.46).

### Question II

We aimed to study the dynamic of sociality at the individual level. First, we analysed the overall effects of sex, age and hierarchical rank on sociality (Model 1.5) and inspected their interaction with *Group*. In a following model (Model 1.6) we included the significant three-way interaction *Group*Sex*Hierarchical rank* but excluded the interaction of *Sex*Age*Hierarchical rank*. We did so in order to reduce the complexity of interpretation and to focus on the different relations between groups rather than the effect of socio-demographic categories on individuals’ sociality.

To deal with the potential effect of genetic link between individuals, we calculated the matri-love of each individual classified as adult females or juveniles (males and females) according to the group they stayed in (Model 2.0). We excluded adult males as they originated from a different natal group. In this model, we kept *Age* and *Sex* as control variables. We included group and hierarchical rank as covariates and individual’s identity as a random intercept. In a following model (Model 2.1), we added the significant interaction *Group*Hierarchical rank* to test for the differential effect of hierarchical rank across groups.

### Question III

In model 3.0, similarly to our analysis of sociality at the group level, the dependent variable was the group grooming reciprocity and the fixed effect was the *Group* with *Year* as random intercept. Finally, in Model 3.1, *Group*, *Sex ratio*, *Age ratio*, *Group size* and the *Time of observation* were used as fixed effects and the *Year* set as a random intercept.

## KEY RESOURCES TABLE

### SUPPLEMENTAL INFORMATION

**Figure S1.**
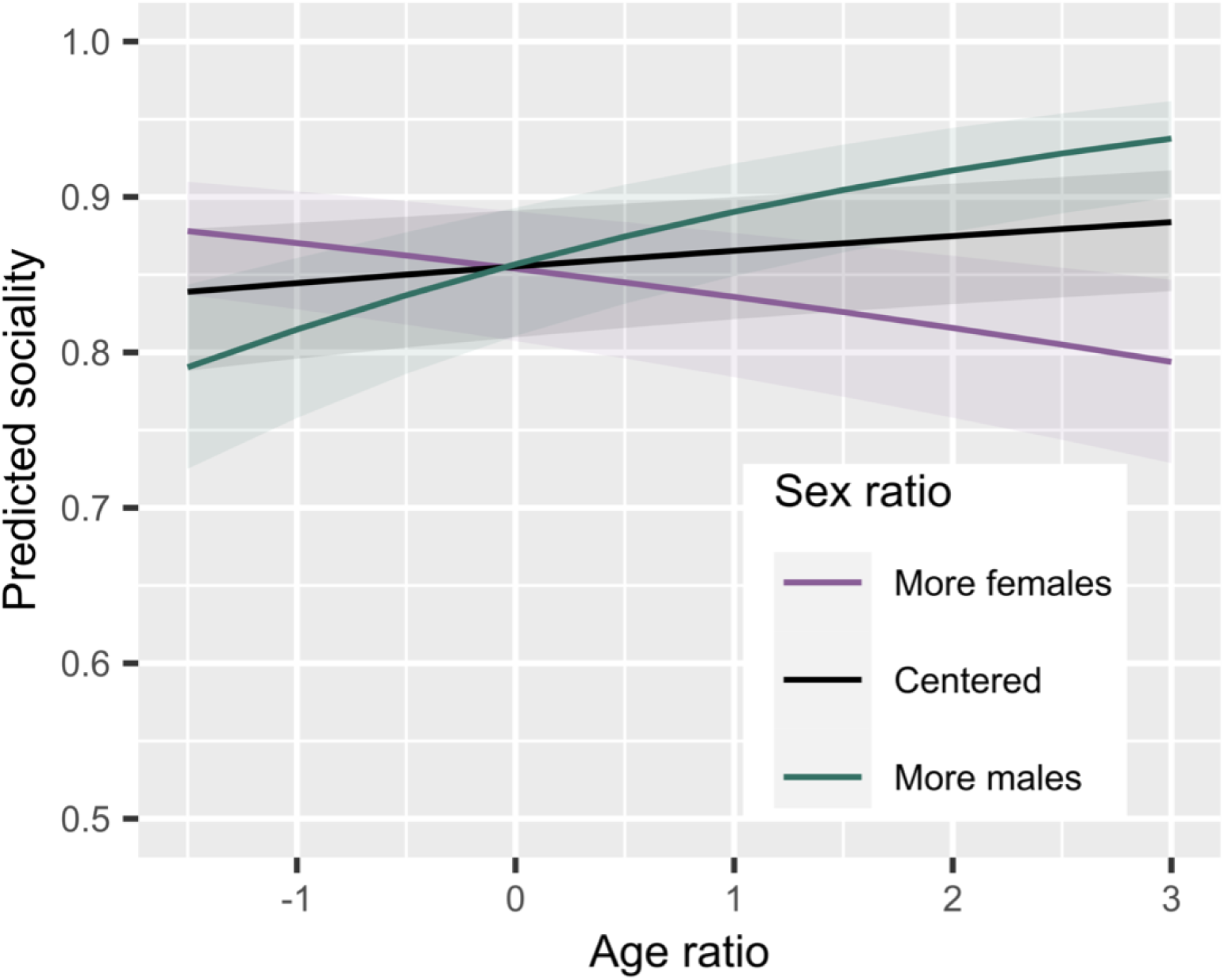
The influence of sex ratio on the effect of age ratio on sociality. *Related to* Figure 1. The x-axis represents the variation in the scaled value of the age ratio, negative values represent a juvenile skewed group and positive values represent an adult skewed group compared to the mean observed age ratio. Shaded areas denote 95% confidence interval (CI).

**Figure S2.**
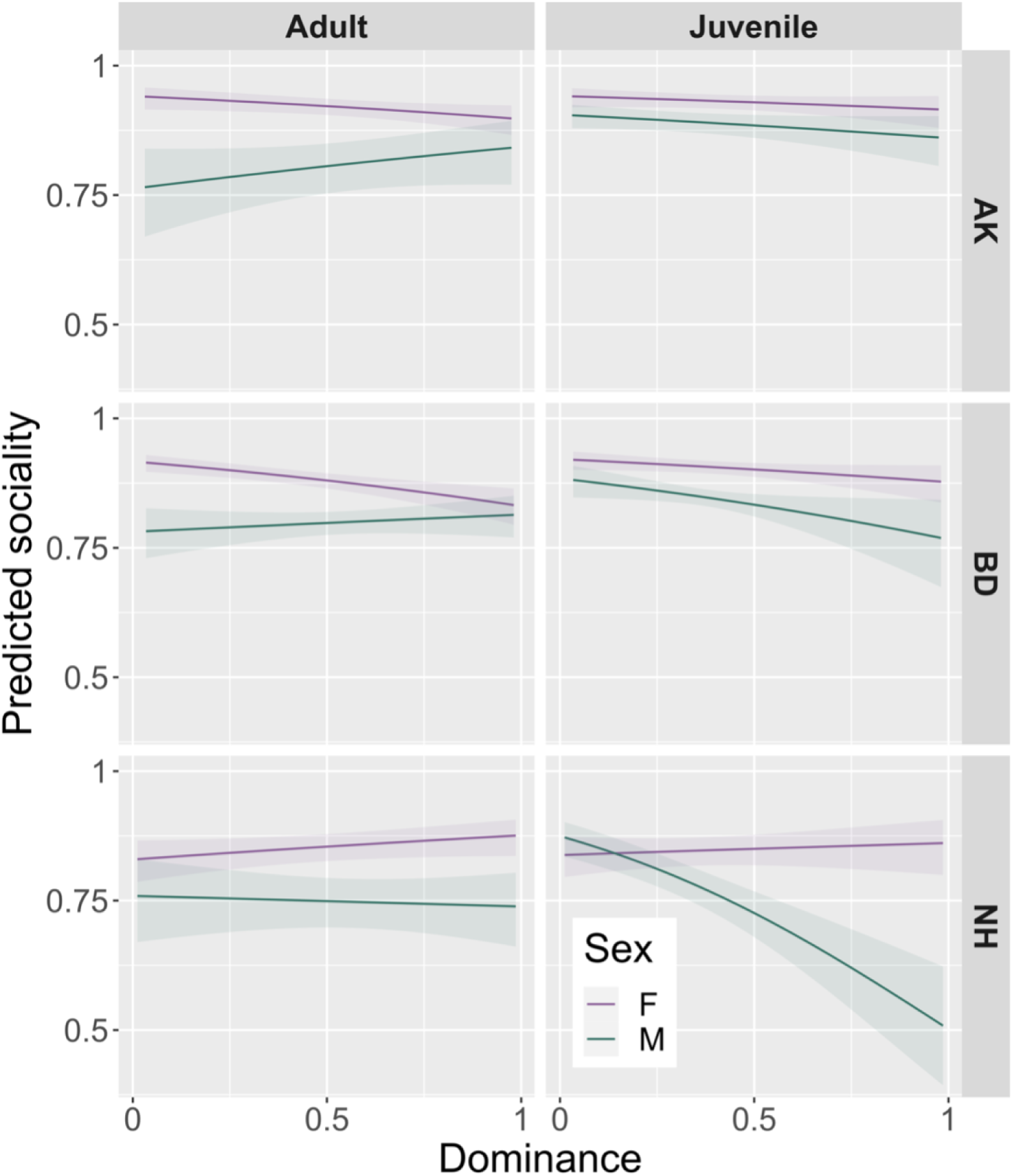
The interaction of sex, age and hierarchical rank on individual sociality within each group. *Related to* Figure 4. Predicted sociality is computed from models predicting the sociality of individuals according to the interaction *Sex* \**Age*\**Hierarchical rank* in each group. Shaded areas denote 95% confidence interval (CI). AK: Ankhase, BD: Baie Dankie, NH: Noha.

**Figure S3.**
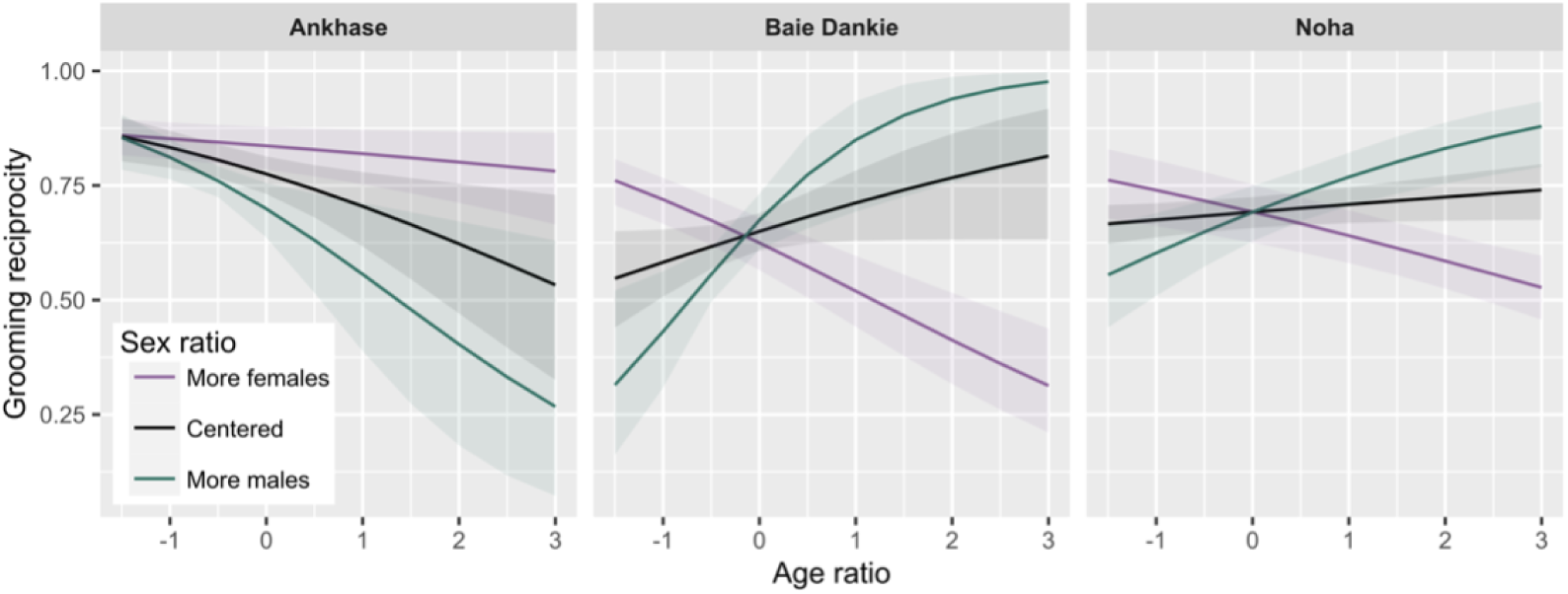
Differences of interaction between age ratio and sex ratio on grooming reciprocity within each group. *Related to* Figure 6. Predicted grooming reciprocity is computed from model predicting the grooming reciprocity according to the interaction *Age-ratio*\**Sex-ratio* in each group. Shaded areas denote 95% confidence interval (CI).

**Table S1.** Ethogram of behaviours included in affiliative interactions. *Related to Methods*.

| Behaviour | Code | Type of interaction | Description |
| --- | --- | --- | --- |
| Embrace | EM/HU | Affiliative | Individual is taking another one in its arm in an affiliative way. |
| Grooming | GR | Affiliative | Individual groom a partner. |
| Infant handling | IH | Affiliative | Individual is manipulating the infant of another individual. |
| Lip smacking | LS | Affiliative | Making noise by clapping lips together, or clicking teeth together. |
| Mouth contact | MM/MC | Affiliative | Individual touches its mouth to/near its partner's (as if giving a kiss). |
| Nursing | NU | Affiliative | Infant is suckling on its mother. |
| Play | PL | Affiliative | Individual play with another. |
| Present | PR | Affiliative | Presenting a part of body to be groomed. |
| Sit in contact | SG | Affiliative | Individuals sitting in contact, one next to another. |
| Sleep with | SW | Affiliative | Individual is sleeping in contact with another individual. |
| Touch | TO | Affiliative | Individual touching a partner in a gentle way. |

**Table S2.** Ethogram of behaviours included in agonistic interactions. *Related to Methods*.

| **Behaviour** | **Code** | **Type of interaction** | **Description** |
| --- | --- | --- | --- |
| Attack | AT | Agonistic (direct) | Forward motion of the body. |
| Bark | BA | Agonistic | Individual produce a specific alarm call type (bark). |
| Bite | BI | Agonistic (direct) | Biting another individual. |
| Chase | CH | Agonistic (direct) | Running after another individual who is fleeing. |
| Crawl | CR | Agonistic | Bow down to an aggressor while looking at them. |
| Fight | FI | Agonistic (direct) | When two or more individuals engage in an aggressive interaction. |
| Flee | FL | Agonistic | Run away from an aggressor as they chase. |
| Grab | GB | Agonistic (direct) | Grabbing another individual. |
| Hit | HI | Agonistic (direct) | Hitting another individual. |
| Jump Aside | JA | Agonistic | Jump aside to avoid something or someone. |
| Kidnapping Baby | KB/SB | Agonistic | Stealing a baby and refusing to hand it back to its mother. |
| Look for support | LO | Agonistic | Looking around for support while they are being threatened by an aggressor. |
| Scream | SC | Agonistic | Distress/scream to an aggressor. |
| Stare | ST | Agonistic | Popping up the eyelids so white above eyes is visible.<br>Often combined with attack. |
| Steal Food | SF | Agonistic | Take another individual's food. |
| Stand Up | SU | Agonistic | Stand up during a conflict with another individual, often following a crawl. |
| Take Place | TP | Agonistic | Individual displaces another and takes their place, usually combined with approach. |

**Table S3.** Ethogram of behaviours included in all types of interactions. *Related to Methods*.

| **Behaviour** | **Code** | **Type of interaction** | **Description** |
| --- | --- | --- | --- |
| Approach | AP | All | Approaching another individual. |
| Auto-groom | AG | All | Individual groom itself. |
| Avoid | AV | All | Head or body movement away from an aggressor, or stopping previous behaviour. |
| Carry infant | CA | All | Individual carries another's infant whilst moving. |
| Feed from mouth | FM | All | Individual takes food from another individual's cheek pouches. |
| Follow | FO | All | Individual clearly walks behind another one. |
| Frustration hop | FH | All | Individual jumps up and down, usually while screaming and potentially looking intensely at another individual. |
| Grunt | GU | All | Individual produces low-frequency guttural calls (grunts). |
| Ignore | IG | All | Carry on with previous behaviour. |
| Leave | LE | All | Walk away. |
| Retrieve baby | RB | All | Mother retrieves her infant from a partner or from the ground if the infant was independent. |
| Retreat | RT | All | Quickly leave the proximity of another individual. |
| Refuse | RF | All | Individual refuses attempts to interact. |
| Smell | SM | All | Individual smells or sniffs its own or another individual |
| Scream | SC | All | Individual produces a distress call/scream usually during agonistic interactions but also produced during frustrating situations. |
| Teeth-chattering | TC | All | jaw movements that are noisy, usually between males of different ranks to initiate close proximity/grooming. |
| Walk by | WB | All | Individual walks parallel (approaches indirectly) to another one. |
| Vocalization | VO | All | Individual produces any call that is not an alarm call or a scream. |

**Table S4.** Demography within each group across year. *Related to Figure 1 and Figure 6*.

| Year | Group | Sex ratio | Age ratio | Group size |
| --- | --- | --- | --- | --- |
| 2012 | Ankhase | 1.31 | 0.58 | 30 |
| 2012 | Baie Dankie | 0.76 | 0.76 | 44 |
| 2012 | Noha | 1.24 | 0.81 | 38 |
| 2013 | Ankhase | 1.57 | 0.50 | 36 |
| 2013 | Baie Dankie | 0.96 | 0.44 | 49 |
| 2013 | Noha | 1.10 | 0.56 | 42 |
| 2014 | Ankhase | 1.33 | 0.67 | 35 |
| 2014 | Baie Dankie | 1.27 | 0.52 | 50 |
| 2014 | Noha | 1.14 | 0.47 | 47 |
| 2015 | Ankhase | 1.33 | 0.45 | 42 |
| 2015 | Baie Dankie | 1.12 | 0.56 | 53 |
| 2015 | Noha | 1.08 | 0.52 | 50 |
| 2016 | Ankhase | 1.20 | 0.47 | 22 |
| 2016 | Baie Dankie | 0.79 | 0.59 | 43 |
| 2016 | Noha | 1.07 | 0.38 | 29 |
| 2017 | Ankhase | 0.64 | 0.64 | 18 |
| 2017 | Baie Dankie | 0.92 | 0.92 | 46 |
| 2017 | Noha | 0.82 | 0.35 | 31 |
| 2018 | Ankhase | 0.46 | 0.36 | 19 |
| 2018 | Baie Dankie | 0.81 | 1.45 | 49 |
| 2018 | Noha | 0.88 | 0.60 | 32 |
| 2019 | Ankhase | 0.86 | 0.53 | 26 |
| 2019 | Baie Dankie | 0.82 | 0.94 | 62 |
| 2019 | Noha | 0.83 | 0.91 | 42 |
| 2020 | Ankhase | 0.82 | 0.82 | 20 |
| 2020 | Baie Dankie | 0.81 | 1.24 | 56 |
| 2020 | Noha | 0.82 | 1.11 | 40 |

**Table S5.** Number of individuals in each socio-demographic category within each group across years. *Related to Figure 4*.

| Year | Group | Adult Females | Adult Males | Juvenile Females | Juvenile Males | Total |
| --- | --- | --- | --- | --- | --- | --- |
| 2012 | Ankhase | 5 | 4 | 5 | 7 | 21 |
| 2012 | Baie Dankie | 14 | 4 | 8 | 4 | 30 |
| 2012 | Noha | 11 | 5 | 4 | 7 | 27 |
| 2013 | Ankhase | 6 | 4 | 5 | 9 | 24 |
| 2013 | Baie Dankie | 11 | 4 | 8 | 12 | 35 |
| 2013 | Noha | 11 | 3 | 6 | 13 | 33 |
| 2014 | Ankhase | 8 | 4 | 4 | 14 | 30 |
| 2014 | Baie Dankie | 10 | 5 | 10 | 20 | 45 |
| 2014 | Noha | 12 | 3 | 7 | 15 | 37 |
| 2015 | Ankhase | 8 | 2 | 5 | 13 | 28 |
| 2015 | Baie Dankie | 12 | 7 | 8 | 15 | 42 |
| 2015 | Noha | 12 | 3 | 8 | 16 | 39 |
| 2016 | Ankhase | 3 | 3 | 6 | 8 | 20 |
| 2016 | Baie Dankie | 12 | 4 | 10 | 13 | 39 |
| 2016 | Noha | 6 | 2 | 7 | 10 | 25 |
| 2017 | Ankhase | 5 | 2 | 4 | 4 | 15 |
| 2017 | Baie Dankie | 15 | 7 | 6 | 15 | 43 |
| 2017 | Noha | 7 | 1 | 7 | 11 | 26 |
| 2018 | Ankhase | 4 | 1 | 6 | 4 | 15 |
| 2018 | Baie Dankie | 17 | 12 | 7 | 6 | 42 |
| 2018 | Noha | 10 | 2 | 6 | 5 | 23 |
| 2019 | Ankhase | 7 | 2 | 5 | 5 | 19 |
| 2019 | Baie Dankie | 20 | 9 | 7 | 7 | 43 |
| 2019 | Noha | 13 | 6 | 4 | 7 | 30 |
| 2020 | Ankhase | 7 | 2 | 4 | 5 | 18 |
| 2020 | Baie Dankie | 20 | 10 | 10 | 12 | 52 |
| 2020 | Noha | 13 | 5 | 8 | 6 | 32 |

